# Reduced STMN2 and pathogenic TDP-43, two hallmarks of ALS, synergize to accelerate motor decline in mice

**DOI:** 10.1101/2024.03.19.585052

**Authors:** Kelsey L. Krus, Ana Morales Benitez, Amy Strickland, Jeffrey Milbrandt, A. Joseph Bloom, Aaron DiAntonio

## Abstract

Pathological TDP-43 loss from the nucleus and cytoplasmic aggregation occurs in almost all cases of ALS and half of frontotemporal dementia patients. *Stathmin2* (*Stmn2)* is a key target of TDP-43 regulation and aberrantly spliced *Stmn2* mRNA is found in patients with ALS, frontotemporal dementia, and Alzheimer’s Disease. STMN2 participates in the axon injury response and its depletion *in vivo* partially replicates ALS-like symptoms including progressive motor deficits and distal NMJ denervation. The interaction between STMN2 loss and TDP-43 dysfunction has not been studied in mice because TDP-43 regulates human but not murine *Stmn2* splicing. Therefore, we generated trans-heterozygous mice that lack one functional copy of *Stmn2* and express one mutant TDP-43^Q331K^ knock-in allele to investigate whether reduced STMN2 function exacerbates TDP-43-dependent pathology. Indeed, we observe synergy between these two alleles, resulting in an early onset, progressive motor deficit. Surprisingly, this behavioral defect is not accompanied by detectable neuropathology in the brain, spinal cord, peripheral nerves or at neuromuscular junctions (NMJs). However, the trans-heterozygous mice exhibit abnormal mitochondrial morphology in their distal axons and NMJs. As both STMN2 and TDP-43 affect mitochondrial dynamics, and neuronal mitochondrial dysfunction is a cardinal feature of many neurodegenerative diseases, this abnormality likely contributes to the observed motor deficit. These findings demonstrate that partial loss of STMN2 significantly exacerbates TDP-43-associated phenotypes, suggesting that STMN2 restoration could ameliorate TDP-43 related disease before the onset of degeneration.

## Introduction

Amyotrophic Lateral Sclerosis (ALS) is a neurodegenerative disease characterized by progressive motor dysfunction and death on average 3-4 years after symptom onset (van Es et al., 2017). ALS is also highly heterogeneous, making it difficult to model and develop treatments. Despite this heterogeneity, one commonality is TDP-43 pathology, which occurs in 97% of ALS patients. TDP-43 is a predominantly nuclear RNA-binding protein that regulates many aspects of RNA processing including splicing and polyadenylation (S. C. Ling et al., 2013). TDP-43 normally functions to maintain healthy levels of correctly processed mRNAs by suppressing both cryptic exon inclusion and use of cryptic polyadenylation sites (Arnold et al., 2024; Bryce-Smith et al., 2024; Jeong et al., 2017; J. P. Ling et al., 2015; Ma et al., 2022; Zeng et al., 2024). In typical ALS, TDP-43 is depleted from the nucleus and mislocalized to the cytosol, where it accumulates in membraneless organelles (Neumann et al., 2006), and causes mitochondrial dysfunction (Wang et al., 2013, 2016, 2017; Yu et al., 2020). Therefore, TDP-43 proteinopathy likely contributes to pathogenesis both through the loss of its normal RNA processing functions and via toxic gain-of-function mechanisms.

Abnormal TDP-43 localization in ALS has been associated with aberrant splicing of many transcripts, foremost *Stathmin2* (*STMN2*). Cryptic exon inclusion in *STMN2* mRNAs leads to dramatically reduced STMN2 protein, both in cellular models and the spinal cords of ALS patients (Klim et al., 2019; Melamed et al., 2019). STMN2, a tubulin-binding protein that regulates microtubule dynamics, is highly enriched in neurons (Morii et al., 2006). Genetic knockdown of the single Drosophila *STMN2* ortholog results in neuromuscular junction (NMJ) instability i.e., presynaptic terminal retraction from the post-synaptic membrane (Graf et al., 2011). In murine sensory neurons, overexpression of stabilized STMN2 delays axon degeneration and maintains axonal mitochondrial motility after injury (Shin et al., 2012). In addition, STMN2 is prominently upregulated in regenerating axons and is required for normal axon regeneration after injury (Klim et al., 2019; Li et al., 2023; Melamed et al., 2019; Shin et al., 2014). To test the hypothesis that STMN2 loss contributes to ALS pathology, we and others generated in vivo mouse models that disrupt STMN2 expression (Guerra San Juan et al., 2022; Krus et al., 2022; López-Erauskin et al., 2024). These studies demonstrated that constitutive STMN2 loss results in severe NMJ pathology that preferentially affects fast-fatigable motor units, as occurs in ALS. Moreover, mice with partially reduced STMN2 expression (*Stmn2*^*+/-*^) exhibit a progressive, late-onset, motor-selective deficit, the hallmark symptom of ALS (Krus et al., 2022). Finally, conditional loss of STMN2 in adult mice triggers motor neuropathy (López-Erauskin et al., 2024). Taken together, these findings support the hypothesis that STMN2 depletion caused by TDP-43 dysfunction contributes to ALS pathology.

TDP-43 regulates the splicing and polyadenylation of many transcripts (Arnold et al., 2024; Bryce-Smith et al., 2024; Jeong et al., 2017; J. P. Ling et al., 2015; Ma et al., 2022; Zeng et al., 2024), but this is unfortunately difficult to model in mice because the cryptic exons TDP-43 regulates are not conserved between humans and mice (J. P. Ling et al., 2015). While this limits the utility of *TARDBP* mutant mice as an ALS model, mice with TDP-43 mutations do exhibit ALS-relevant phenotypes, likely due to toxic *gain-of-function* mechanisms, or loss of other better conserved TDP-43 functions. In this study, we test the hypothesis that ALS-like pathology is potentiated by the combination of TDP-43 toxic gain-of-function and a key consequence of TDP-43 *loss*-of-function, decreased STMN2. To achieve this, we generated mice that lack one copy of *Stmn2* (*Stmn2*^+/-^) and carry one copy of an endogenous TDP-43 allele equivalent to a known pathogenic variant (Q331K) found in ALS patients (Sreedharan et al., 2008). TDP-43^Q331K/Q331K^ homozygous mice display ALS-relevant phenotypes including altered splicing of murine target mRNAs, progressive weakness and NMJ defects (White et al., 2018). Cellular models of this mutation also reveal increased DNA damage and defects in mitochondrial morphology and motility (Guerrero et al., 2019; Wang et al., 2013). However, nuclear TDP-43 levels are not reported to decrease in this model. In fact, TDP-43 protein is elevated due to disrupted autoregulation (White et al., 2018). Thus, the point mutation likely mediates gain-of-function effects associated with TDP-43 proteinopathy.

Excitingly, TDP-43^Q331K/+^; *Stmn2*^*+/-*^ trans-heterozygous mice exhibit an enhanced motor deficit compared to either single mutant alone, and without obvious sensory or coordination defects. Surprisingly, this motor deficit occurs without motor neuron loss, axon degeneration, or NMJ denervation. Instead, these two alleles have a combined deleterious effect on distal mitochondria. Interestingly, prior studies of STMN2 and TDP-43 demonstrate that each facilitates mitochondrial motility and that mutant TDP-43 inhibits mitochondrial fusion and fission (Shin et al., 2012; Wang et al., 2013). Our findings suggest that even partial loss of STMN2 may exacerbate the dysregulation of mitochondrial homeostasis by mutant TDP-43. Thus, while TDP-43 has many molecular targets, restoration of STMN2 expression alone may be sufficient to significantly ameliorate TDP-43-dependent pathology, a strategy currently being tested in clinical trials (National Institute of Health, 2022).

## Methods

### Inverted Screen Assay

Mice were placed on a wire mesh screen and the screen was inverted. Each mouse underwent 3 trials with five-minute intervening rest periods. Latency to fall for each mouse was recorded. The maximum time allowed for the task was 2 minutes. The average of the three trials was taken.

### Rotarod Assay

Mice were trained the day prior to testing by walking on the rotarod (Panlab, LE8205) at constant speed for five minutes. On the day of testing, they were trained for another five minutes. Each mouse then underwent five trials on the rotarod that started at 2 rpm and accelerated 1 rpm every three seconds to a maximum of 24 rpm. They were allowed to rest for five-minutes between trials. Latency to fall off the rod was recorded and the average of the five trials was taken.

### Von Frey Assay

The Von Frey assay was performed according to the up-down method (Chaplan et al., 1994). Briefly, mice were individually habituated on a wire mesh screen in plexiglass boxes with opaque dividers between mice. They stayed in the boxes for four hours to acclimate before testing. The hind paw plantar surface was stimulated for 2 seconds with a filament, beginning with a force of 0.32 g. If the mouse withdrew the paw, the next filament used would be one that delivered less force. A filament that delivered greater force would be tested if withdraw did not occur. Stimulation would then either decrease or increase in force until the mouse ceased or began withdrawing, respectively. Stimulation occurred four more times after that first change in response. The 50% withdraw threshold was calculated (Dixon, 1980). Results from right and left hind paws were averaged for each mouse. Mice were always tested at the same time of day and were given a rest period between trials.

### Nerve Electrophysiology

Compound muscle action potentials (CMAPs) were acquired using a Viking Quest electromyography device (Nicolet) as previously described (Beirowski et al., 2011). Briefly, mice were anesthetized, a recording electrode placed in the plantar surface of the mouse’s foot and a stimulating electrode in the sciatic notch. Supramaximal stimulation was used for CMAPs. SNAPs were acquired using the same device, as previously described (Geisler et al., 2016). Mice were anesthetized and electrodes placed subcutaneously with the stimulating electrode placed in the tail tip 30 mm distal from the recording electrode placed in the base of the tail. A ground electrode was placed in between. Supramaximal stimulation was used for SNAPs.

### Analysis of Brains and Spinal Cords

Mice were intracardially perfused, first with PBS, then 4% PFA. Spinal columns and brains were further fixed in 4% PFA for 3 days after dissection. Spinal cords were then dissected out and both tissues transferred to 30% sucrose until no longer buoyant. Spinal cords were divided into cervical, thoracic, and lumbar segments and the tissues were embedded in OCT and cut into 20 µm thick sections by cryotome. Slides were place on a 55°C hot plate for one hour, then left to dry overnight at RT. Slides were stored at -20°C until use. Cresyl Violet & Luxol Blue staining of brain sections was performed using a commercially available kit according to the protocol provided (Abcam, ab150675). Brain images were taken on a Zeiss Axio Scan Z1 slide scanner.

Antigen retrieval was used to detect ChAT, TDP-43, and vGlut. Spinal cord sections on slides were first put in a plastic or glass Coplin jar with 1x citrate buffer solution at pH 6.0 (Sigma C9999), then boiled in a pressure cooker for 2 min. Including the time required to arrive at temperature, boil for two minutes, and to release pressure, slides remain in the pressure cooker for about 30 minutes. Slides are then left to incubate another 30 minutes in the warmed 1x citrate solution and rinsed with tap water to remove the citrate buffer. Following antigen retrieval, slides are washed 2x 5 minutes in PBS, incubated in 0.1% Triton-X in PBS for 30 minutes and blocked in 4% BSA + 1% Triton-X in PBS for 30 minutes. Primary antibody is diluted in blocking buffer and slides are incubated in primary antibody overnight at 4°C (ChAT, Millipore, AB144P, goat, 1:200; vGlut1, EMD Millipore, AB5905, guinea pig, 1:300; TDP-43, Proteintech, 1:1000). Primary antibody is removed the next day and slides washed in PBS + 0.1% Triton 3x 5 minutes. Secondary antibodies (1:250 anti-goat and anti-guinea pig Jackson Immunoresearch, Cat # 705-166-14 and 706-165-148) are diluted in wash solution and slides are incubated 3 hours at RT, then washed 3x 5 minutes in PBS. Vectashield with DAPI is added to sections and slides are protected with a coverslip. Z-stack images were acquired on a Zeiss confocal and maximum projection was utilized to produce images. For GFAP (1:1000, Agilent Technologies, Z033429-2), Iba1 (1:500, Fujifilm, 019-19741), and CD68 (1:200, Bio-Rad, MCA1957GA) staining, the protocol was followed omitting the antigen retrieval and starting at the initial PBS washes. All Images were analyzed using ImageJ.

### Western Blot

Sciatic nerves dissected from mice were immediately flash frozen and kept on dry ice. RIPA buffer with protease inhibitor was added to the tube. Lysis was performed by sonification, and the lysates stored on ice. Sciatic nerve lysates were resolved using SDS polyacrylamide gel electrophoresis (PAGE) on 4-20% Mini-Protean Gel (BioRad), followed by immunoblotting for STMN2 (1:1000 anti-SCG10 (Shin et al., 2012)) and TUJ1 (anti-B3 tubulin, 1:10000 Sigma-Aldrich, T2200), and visualized using standard chemiluminescence. Band intensity was quantified using ImageJ. Blots were normalized to their respective loading controls and then normalized to the control sample.

### Nerve structural analysis

Femoral nerves were processed as previously described (Geisler et al., 2016). Briefly, nerves were fixed in 3% glutaraldehyde in 0.1 ml PBS overnight at 4°C. They were then washed and stained with 1% osmium tetroxide (Sigma Aldrich) overnight at 4°C. The following day, nerves were washed and dehydrated in a serial gradient of 50% to 100% ethanol. Then they were incubated in 50% propylene oxide/50% ethanol, then 100% propylene oxide. Nerves were incubated in Araldite resin/propylene oxide solutions in 50:50, 70:30, 90:10 ratios for 24 hours each, and subsequently embedded in 100% Araldite resin solution (Araldite: DDSA: DMP30, 12:9:1, Electron Microscopy Sciences) and baked overnight at 60°C. Semithin 400—600 nm sections were cut using a Leica EM UC7 Ultramicrotome, placed on microscopy slides, and stained with Toluidine blue. Staining and quantification were performed as previously described (Sasaki et al., 2018). All quantification was performed blinded.

### Tibial Nerve Staining

Tibial nerves were collected and fixed for 1 hr in 4% PFA, then transferred to 30% sucrose overnight at 4 degrees Celsius. Nerves were then embedded in OCT and sectioned on a cryotome at 6 µm. Slides were dried overnight. The slides were fixed in -20°C acetone for 10 minutes, air dried, washed in PBS 2x 5 minutes, and then washed in PBS + 0.1% Triton-X for 10 minutes. Slides were then blocked in 4% BSA + 1% Triton-X for 30 minutes. CD68 primary antibody (1:200, Bio-Rad, MCA1957GA) was diluted in blocking buffer and solution was left on the slides overnight at 4°C. The following morning, the slides were washed 3x 5 minutes in PBS, then incubated in PBS + 0.1% Triton and secondary antibody (Invitrogen, 1:500) for 1 hour at room temperature. The slides were washed 3x 5 minutes in PBS. Vectashield with DAPI was applied to the slides and coverslips were put on. Z-stack images on a Zeiss confocal were taken and maximum projection images were produced. Images were analyzed with ImageJ.

### NMJ analysis

Lumbrical muscles were collected from animals perfused with PBS then 4% PFA. Dissected muscles were incubated in 2% Triton-X in PBS for 30 minutes and blocked with 5% BSA 1% Triton-X for 30 minutes. They were incubated at 4°C overnight in blocking buffer with antibodies against neurofilament and SV2 (2H3, 1:100, DHSB AB2314897; SV2, 1:200, DSHB AB2315387). The following day, muscles were washed in PBS 3x 15 min, then in 1% Triton-X in PBS with FITC rabbit anti-mouse IgG1 (1:400, Invitrogen A21121) and Alexa fluoro-568 conjugated α-bungarotoxin (1:500, Biotium 00006) at room temperature for 3 hours. Muscles were washed 3 x 15 minutes in PBS and whole mounted on slides with Vectashield Mounting medium.

To analyze NMJ morphology, z-stack images of mounted lumbricals were obtained using a confocal microscope. Maximal projection was applied. Individual NMJs were categorized as 1) fully innervated, 2) partially innervated, or 3) not innervated. Over 50 NMJs were evaluated per animal. Images shown are representative images taken with a 40x oil immersion lens. Researchers were blinded to genotype during imaging and image analysis.

### Transmission Electron Microscopy

Mice were perfused with PBS then 4% PFA. Whole feet were removed from the animal and immersed in 3% glutaraldehyde for 3 days. Lumbrical muscles were dissected out, rinsed with 0.1M phosphate buffer and placed in 1% osmium tetroxide at 4°C overnight. They were subsequently washed with 0.1M phosphate buffer 3 times and dehydrated. Dehydration consisted of 30-minute incubations with increasing concentrations of acetone in water: 50%, 70%, 90%, and finally three times in 100% acetone. Muscles were then incubated in Spurr: acetone at ratios of 1:2, 1:1, 3:1 and finally 100% Spurr, with each incubation lasting 24 hours at room temperature (Spurr Resin kit, Electron Microscopy Science, 14300). Lumbricals were embedded in 100% Spurr and polymerized at 65°C for 48 hours. Thin sections were cut by the Washington University Core for Cellular Imaging (WUCCI) and placed on screens for imaging. A JEOL JEM-1400 Plus transmission electron microscope was used for imaging. 6-10 images were taken per sample. Presynapse mitochondrial circularity was quantified using the formula Circularity = 4π*area/perimeter^2^, as described (Abbade et al., 2020). This value approaches 0.0 for increasingly elongated mitochondria, while a perfect circle has a circularity value of 1.0. Synaptic vesicle density was calculated by counting the number of synaptic vesicles in a representative region of the presynapse (without other interfering structures) divided by the area of the region. Cristae clearing was calculated by using the mean grey value of each mitochondrion normalized to the average grey value for all wildtype mitochondria. In this case, a larger value indicates more area of mitochondria not covered by dense cristae. Only mitochondria and synaptic vesicles from intact axons were quantified. The researcher was blinded to genotype during image analysis.

## Results

### Combining pathogenic TDP-43 and reduced STMN2 accelerates progressive motor decline

To investigate the potential synergy between the STMN2 depletion associated with TDP-43 *loss-of-function* and TDP-43 *gain-of-function* toxicity, we crossed *Stmn2*^*+/-*^ and TDP-43^Q331K/+^ mice to generate *TDP-43*^*Q331K/+*^;*Stmn2*^*+/-*^ trans-heterozygous (TS) mice. We hypothesized that by combining these two important features of TDP-43 dysfunction, these mice would surpass prior murine models in recapitulating ALS phenotypes. To ensure that the TDP-43 mutation itself does not alter STMN2 levels, we first harvested sciatic nerves from wild-type, *TDP-43*^*Q331K/+*^, *Stmn2*^*+/-*^, and TS mice (Fig 1A, B) and detected STMN2 by Western blot. As expected, *Stmn2*^*+/-*^ and TS mice show a ∼50% reduction in STMN2 compared to WT or *TDP-43*^*Q331K/+*^, confirming that the mutant TDP-43 allele does not influence STMN2 levels in mice. Thus, we may attribute phenotypic differences between *Stmn2*^*+/-*^ and TS mice to interactions between the two defects rather than a further decrease of STMN2.

**Figure 1:**
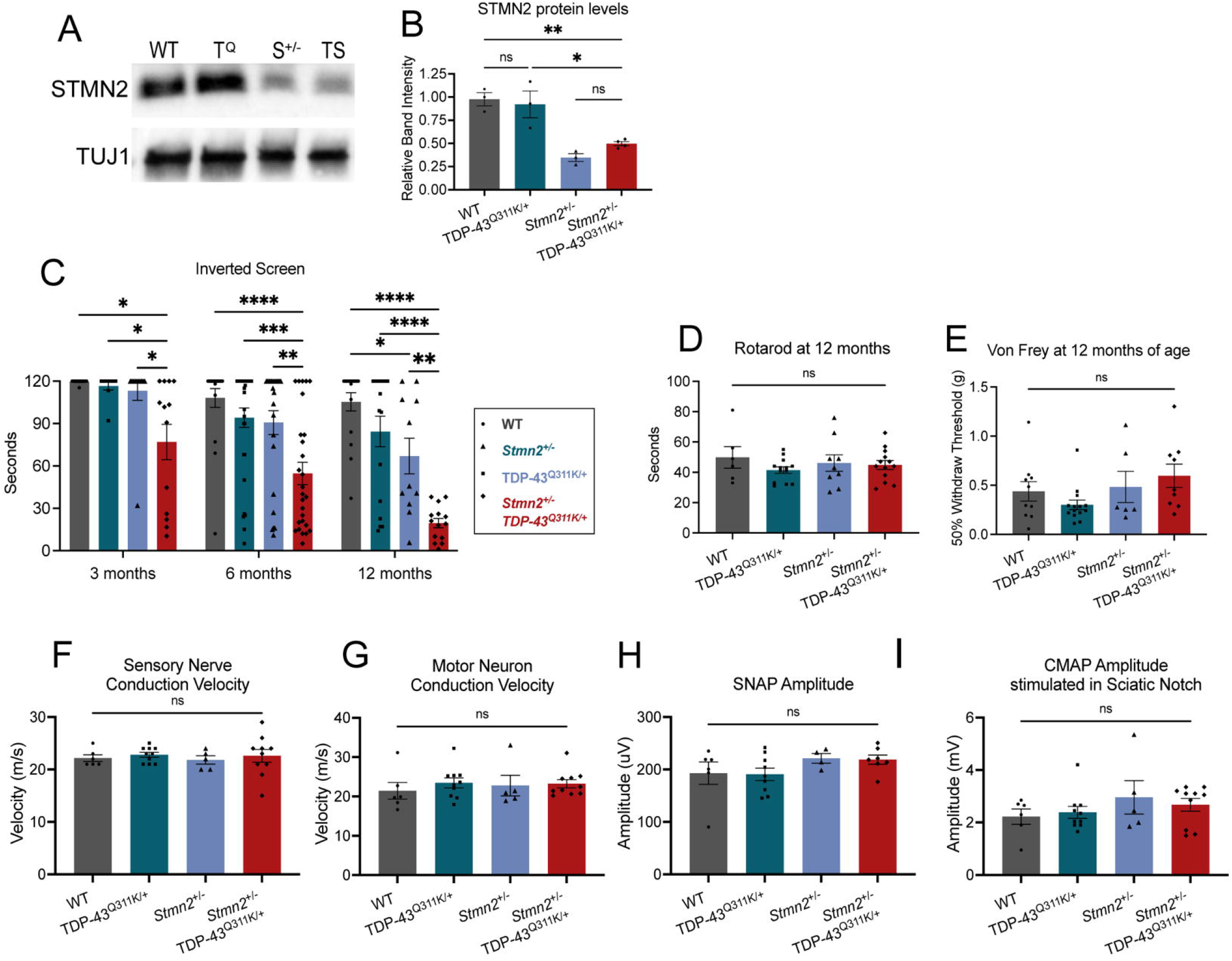
TDP-43^Q331K^; *Stmn2*^*+/-*^ mice have progressive, motor-selective behavioral deficits. **A)** Representative Western blot of STMN2 protein from sciatic nerves of 6-month-old wild type (WT), TDP-43^Q331K^ (T^Q^), *Stmn2*^*+/-*^ (S^+/-^), and TDP-43^Q331K^;*Stmn2*^*+/-*^ (TS) mice. **B)** Quantification of STMN2 protein for each genotype. **C)** Latency time to fall from an inverted screen (max. 120 sec). Statistical significance is determined using 2-way ANOVA with Tukey’s multiple comparison test. **D)** Time (seconds) on an accelerating rotarod before falling, and **E)** average 50% hind paw withdrawal threshold when force (grams) applied, for 12-month-old WT, TDP-43^Q331K^, *Stmn2*^*+/-*^, and TDP-43^Q331K^;*Stmn2*^*+/-*^ mice. **F)** Sensory nerve and **G)** motor nerve conduction velocity. **H)** Sensory nerve action potential amplitude and **I)** compound muscle action potential (CMAP) amplitude stimulated in the sciatic notch for 6-month-old WT, TDP-43^Q331K^, *Stmn2*^*+/-*^, and TDP-43^Q331K^;*Stmn2*^*+/-*^ mice. Unless otherwise stated, statistical significance was determined by ordinary one-way ANOVA with Tukey’s multiple comparisons test. (ns: not significant, *p<0.05, **p<0.01, ***p<0.001, ****p<0.0001).

To assess motor behavior, we tested muscle strength using the inverted screen assay at three, six, and twelve months of age. The TDP-43^Q331K/+^; *Stmn2*^*+/-*^ TS mice exhibit a progressive motor deficit already evident at three months when both TDP-43^Q331K/+^ and *Stmn2*^*+/-*^ mice are indistinguishable from WT mice. As reported, TDP-43^Q331K/+^ mice do not develop motor deficits within 12 months (White et al., 2018), while *Stmn2*^*+/-*^ mice develop late-onset motor deficits (Krus et al., 2022) but still perform significantly better than TS mice at 12 months (Figure 1C). We also used rotarod and Von Frey tests to assay coordination and sensory acuity but detected no significant differences among the four genotypes (Figure 1D, E), demonstrating the motor-selectivity of the TS phenotype.

We next performed electrophysiological analysis of motor and sensory fibers by assaying sensory nerve action potentials (SNAP) and compound muscle action potentials (CMAP). These studies were performed at 6 months of age, when the largest difference in the inverted screen test was observed between the TS mice and each single heterozygote. No significant differences in nerve conduction velocities, SNAP amplitude, or CMAP amplitude were observed across genotypes (Figure 1F-I). These findings suggest that myelination and axon number are likely normal in both sensory and motor fibers.

### Motor axons and NMJs are intact in TDP-43^Q331K/+^;*Stmn2*^+/-^ mice

It was surprising that six-month-old TS mice have normal compound muscle action potentials despite their poor motor performance (Figure 1G). Therefore, to further examine the source of their motor deficit, we performed neuropathological analyses on 6-month-old animals. We first quantified axon density in the predominantly-motor femoral nerve. Consistent with their normal electrophysiology, we found no axon loss in TS animals (Fig 2A,B). Next, we used CD68, a marker of activated macrophages, to search for evidence of neuroinflammation in the TS animals that could contribute to the motor dysfunction, however we found no significant increase of CD68+ macrophages in their tibial nerves (a distal motor-dominant nerve) (Fig 2C,D). We next examined their lumbrical muscles for evidence of denervation by immunohistochemically labeling the pre- and post-synaptic structures at their neuromuscular junctions (NMJs). Lumbricals are distal, predominantly fast-twitch muscles and therefore especially vulnerable to ALS pathology. However, we did not find significant lumbrical muscle denervation in the TS mice. Rather, we observed abundant innervating axons that were well apposed to postsynaptic endplates (Fig 2E, F). Hence, the progressive motor deficit observed in TS mice is not explained by either axon or NMJ loss.

**Figure 2:**
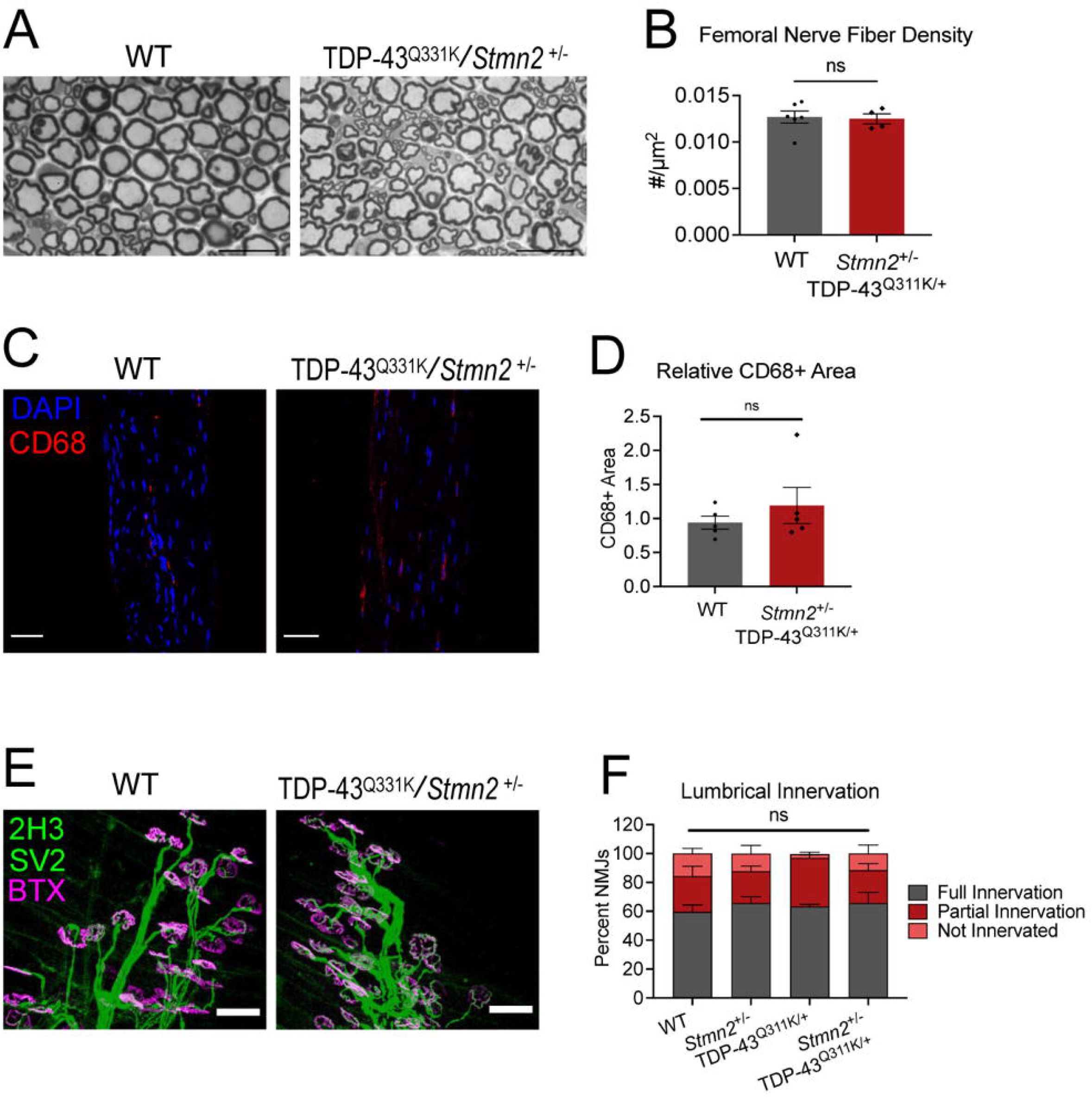
TDP-43^Q331K^; *Stmn2*^*+/-*^ mice do not have obvious degeneration in the peripheral nervous system. **A)** Toludine blue staining of femoral nerves from 12-month-old WT and TDP-43^Q331K^;*Stmn2*^*+/-*^ mice. **B)** Quantification of femoral nerve axon density. Statistical significance determined by Student’s t-test. **C)** Representative images of 6-month-old WT and TDP-43^Q331K^;*Stmn2*^*+/-*^ tibial nerve sections stained with anti-CD68 to detect macrophages. Scale bar = 50 µm. **D)** Quantification of macrophage number/field. Statistical significance determined by Student’s t-test. **E)** Representative images of lumbrical muscle NMJs labelled with anti-neurofilament and anti-SV2 (synaptic vesicles) in green and fluorescently labelled bungarotoxin marking endplates in magenta. Scale bar = 50 µm. **F)** Quantification of lumbrical muscle innervation. Statistical significance determined using a 2-way ANOVA with Sidak’s multiple comparisons test. (ns: not significant)

### The CNS does not show obvious signs of degeneration in TDP-43^Q331K^;*Stmn2*^+/-^ mice

To investigate whether motor deficits in TS mice could be caused by cortical neuron degeneration, we stained brains from adult WT and TS mice with cresyl violet to measure the cortical thickness of the primary and secondary motor cortices. TS animals did not show signs of cortical thinning or neuron loss compared to WT animals. Both TDP-43 dysfunction and STMN2 depletion can affect axon outgrowth (Fallini C, 2012; Schmid B, 2013; Morii H, 2006; Klim JR, 2019; Melamed Z, 2019), so we assessed cortical white matter by counterstaining these sections with luxol blue, a dye that binds myelin. Corpus callosum thickness was not significantly different in TS and WT animals. Thus, overall cellular and axonal structure appears to be maintained in the CNS of TS mice (Fig 3A).

**Figure 3:**
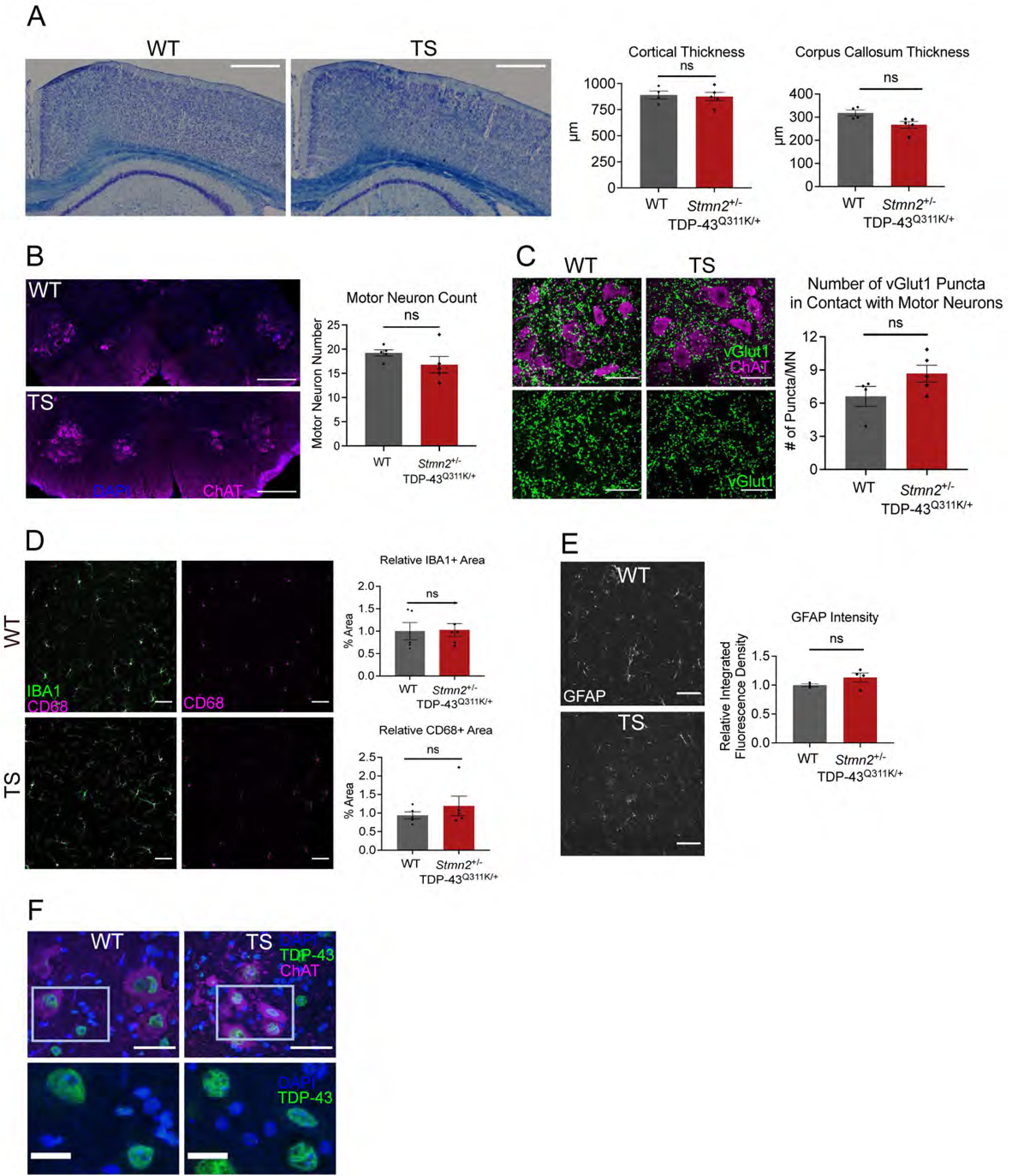
TDP-43^Q331K^; *Stmn2*^*+/-*^ mice do not have obvious degeneration in the central nervous system. **A)** Representative images of 6-month-old WT and TDP-43^Q331K^;*Stmn2*^*+/-*^ coronal brain sections stained with luxol blue (myelin) and cresyl violet (cell bodies), quantified on the right. Scale bar = 100 µm. **B)** Motor neurons in lumbar spinal cord sections labelled with anti-ChAT in magenta, quantified on the right. (ns: not significant) **C)** Representative images of spinal cord sections labelled for motor neurons (ChAT) in magenta and upper motor neuron pre-synapses (vGlut1) in green. Scale bar = 50 µm. Quantification on the right. **D)** Representative images of spinal cord sections labelled for microglia with anti-IBA1 in green and anti-CD68 antibodies in magenta. Scale bar = 50 µm. Quantification below. **E)** Representative images of spinal cord sections labelled for astrocytes with anti-GFAP. Scale bar = 50 µm. Quantification to the right. Statistical significance determined using Student’s t-test. (ns: not significant) **F)** Representative images of TDP-43 localization in spinal cord sections labelled with DAPI in blue, anti-TDP-43 in green, and anti-ChAT in magenta. Scale bar top = 50 µm, scale bar bottom = 10 µm.

We next assessed degeneration in the lumbar spinal cord, where motor neurons innervating the distal hindlimbs are located. Quantifying choline acetyltransferase (ChAT)-positive motor neurons showed that TS mice had the same number of motor neurons per spinal cord section as WT animals, indicating that motor neuron death was not responsible for the behavioral deficits (Fig 3B). Another feature of ALS is the loss of spinal cord synaptic integrity (Broadhead et al., 2022). To examine the connection between upper and lower motor neurons, we stained for vGlut1, a marker of upper motor neuron presynapses, and ChAT, the lower motor neuron marker, and counted the number of vGlut1-positive puncta adjacent to spinal motor neurons. However, we found no significant difference in vGlut1 puncta per motor neuron in TS spinal cords, indicating that innervation of spinal motor neurons by upper motor neurons is intact in these mice (Fig 3C). Next, we assessed inflammatory changes in the spinal cord, as inflammation and gliosis are prominent features in ALS patients and ALS mouse models (Bright et al., 2021). We used antibodies against IBA1 to label microglia, CD68 to label activated microglia, and GFAP to label activated astrocytes, and found no significant difference in microglia area, reactive microglia abundance, or astrocytosis in the spinal cords of TS mice (Fig 3D, E). Hence, we find no evidence of reactive inflammation that could explain their motor deficits. Finally, we examined whether STMN2 depletion induces mislocalization of the mutant TDP-43 to the cytosol where it could have multiple deleterious effects. However, TDP-43 protein appears tightly localized to the nucleus in both WT and TS spinal cord (Fig 3F). Taken together, these data suggest that motor deficits in TS mice are unlikely to be explained by brain or spinal cord pathology.

### TDP-43^Q331K^; *Stmn2*^*+/-*^ mice have mitochondrial defects in distal axons and NMJs

Finding the gross neuronal structure of TS mice intact, we next performed an ultrastructural analysis of their distal axons and neuromuscular junctions. We prepared lumbrical muscles from the hind paw for transmission electron microscopy (TEM) and examined mitochondria in the nerves penetrating these muscles, as well as the mitochondria and synaptic vesicles within the NMJs. We assessed the circularity of the mitochondria and observed that mitochondria were significantly rounder in both the distal axon and NMJ of TS mice compared to TDP-43^Q331K/+^, *Stmn2*^*+/-*^, or WT mice (Fig 4A-C) suggesting an imbalance in mitochondrial fission and fusion. We did not observe a significant difference in mitochondrial size (Sup fig 1A). In addition, there was a tendency for mitochondria in the TS to exhibit less dense cristae, the site of electron transport chain activity (Sup Fig 1B). Finally, we quantified synaptic vesicle density across genotypes but did not observe a statistically significant difference (Fig 4D). In summary, TDP-43^Q331K/+^ /*Stmn2*^*+/-*^ mice display altered mitochondria morphology in their distal axons and NMJs that likely reflect defective mitochondrial dynamics and dysfunction that may explain the motor deficits present in these mice.

**Figure 4:**
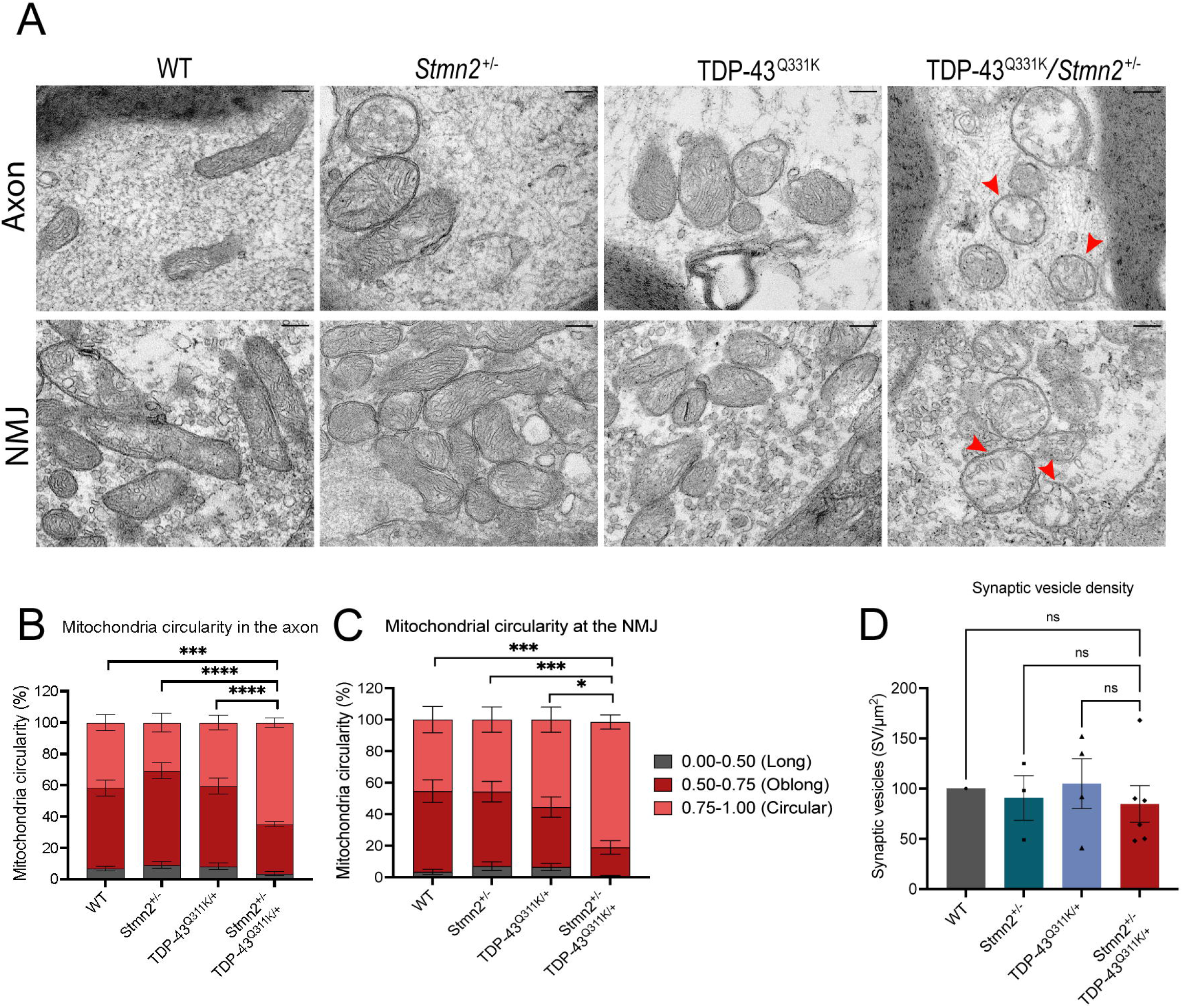
TDP-43^Q331K^; *Stmn2*^*+/-*^ mice have mitochondrial abnormalities in distal nerves. **A)** Representative transmission electron microscopy images from distal nerves of WT, TDP-43^Q331K^, *Stmn2*^*+/-*^, and TDP-43^Q331K^;*Stmn2*^*+/-*^ mice. Arrows mark round mitochondria. Scale bar = 200 nm. **B**,**C)** Quantification of mitochondrial circularity in the distal axon **(B)** and NMJ **(C)**. Statistical significance determined by 2-way ANOVA with Tukey’s multiple comparisons test. **D)** Quantification of NMJ synaptic vesicle density. Statistical significance determined by ordinary one-way ANOVA with Tukey’s multiple comparisons. (ns: not significant, **p<0.01, ***p<0.001, ****p<0.0001)

## Conclusions

*STMN2* has been consistently identified as the most reduced transcript associated with TDP-43 dysfunction in human cellular models of ALS (Klim et al., 2019; Melamed et al., 2019). STMN2 protein has functions in both axon degeneration and regeneration (Klim et al., 2019; Melamed et al., 2019; Shin et al., 2012, 2014) leading to the hypothesis that STMN2 depletion partly mediates the pathological consequences of TDP-43 dysfunction. However, TDP-43 does not regulate *Stmn2* splicing in mice, making it difficult to assess the effects of STMN2 loss in the setting of TDP-43 dysfunction in murine models. A recent study developed a mouse carrying a humanized *Stmn2* gene to test this hypothesis. While the model was useful to study *Stmn2* splicing *in vivo*, it was unsuccessful at assessing pathological enhancement because the TDP-43^Q331K^ mutant mouse used did not feature TDP-43 mislocalization from the nucleus, and thus STMN2 expression was unaffected (Baughn et al., 2023). Therefore, another approach was required. To that end, we generated a mouse heterozygous for both a *Stmn2* deletion and the TDP-43^Q331K^ allele to directly decrease STMN2 protein levels in the presence of mutant TDP-43. We chose the heterozygous TDP-43^Q331K^ knock-in allele because we expected that its relatively mild pathogenicity would enable detection of synergism between the two alleles. Gratifyingly, we did observe such synergy leading to early onset, motor-selective progressive dysfunction. However, this behavioral phenotype is not accompanied by detectable neuropathology in the brain, spinal cord, peripheral nerves or NMJs. However, their axonal and synaptic mitochondrial morphology is altered, with a preponderance of round mitochondria consistent with dysfunction and an imbalance of mitochondrial fission/fusion. As both STMN2 and TDP-43 influence mitochondrial dynamics, and neuronal mitochondrial dysfunction is a key contributor to neurodegenerative disease (Jadiya et al., 2021), this is a plausible cellular locus for the behavioral deficit.

The identification of transcripts depleted by TDP-43 dysfunction suggested the restoration of those transcripts as a potential treatment for ALS and FTD. Indeed, clinical trials are underway with an ASO intended to restore proper splicing, and hence protein expression, of STMN2 (National Institute of Health, 2022). However, as many different transcripts are altered by TDP-43 dysfunction, it is unclear whether restoring any single target would be sufficient to impact disease pathology, and targeting multiple transcripts simultaneously may be unrealistic. While a definitive answer awaits the outcome of clinical trials, we posit that restoration of normal STMN2 protein levels will significantly ameliorate ALS pathology. First, a partial reduction of STMN2 in the absence of TDP-43 dysfunction does induce a slowly progressive motor neuropathy in mice (Krus et al., 2022), and importantly, this phenotype also arises when STMN2 is depleted post-development in *adult* mice (López-Erauskin et al., 2024). Second, here we show that only a 50% reduction of STMN2 in the background of mildly pathogenic TDP-43 is sufficient to accelerate motor decline, leading to deficits evident by only three months of age, far earlier than in either single mutant model alone. In this scenario, restoration of STMN2 would be predicted to restore function to that of the TDP-43^Q331K/+^ mouse, which shows no behavioral defects even at one year of age. Thus, full restoration of STMN2 would significantly delay disease onset. While there are clearly many important loss- and gain-of-function phenotypes associated with TDP-43 dysfunction that contribute to neurodegeneration in ALS, FTD, and AD, these findings support the continued efforts to target STMN2 as a treatment for these devastating diseases.

## Acknowledgments

We would like to thank members of the DiAntonio and Milbrandt labs for their thoughtful discussions on the study. We would also like to thank Cassidy Menendez, Rachel McClarney, Alicia Neiner, Sylvia Johnson, Xiaolu Sun, Kelli Simburger, Yo Sasaki, and Liya Yuan for their technical support and/or technical training and advice. We also thank the Washington University Core for Cellular Imaging (WUCCI) for their technical support, expertise, and training on the spinning disk microscope and transmission electron microscope and the Genome Engineering & Stem Cell Center (GESC@MGI) at Washington University for generating the *Stmn2* mice.

## Author Contributions

KLK, JM, AJB and AD and conceived of the overall study. KLK collected mouse behavior data and performed TEM and interpreted electron micrographs. KLK, AS and AMB collected mouse tissue samples and performed immunofluorescent staining. AS performed electrophysiology studies. KLK and AMB performed confocal imaging and image analysis. KLK wrote the manuscript and KLK and AMB prepared the figures. AJB, AD, and JM oversaw the analysis and revised the manuscript. All authors gave final approval of the manuscript.

## Figure Legends

**Supplemental Figure 1: TDP-43**^**Q331K**^**; *Stmn2***^***+/-***^ **mice have reduced mitochondrial cristae in distal nerves**

**A)** Average mitochondrial area in distal axons and NMJs in lumbrical muscles of WT, TDP-43^Q331K^, *Stmn2*^*+/-*^, and TDP-43^Q331K^;*Stmn2*^*+/-*^ mice. Each point represents the average area of multiple mitochondria from a single animal. **B)** Mitochondrial cristae density in distal axons and NMJs in lumbrical muscles of WT, TDP-43^Q331K^, *Stmn2*^*+/-*^, and TDP-43^Q331K^;*Stmn2*^*+/-*^ mice calculated using the mean grey value of each mitochondrion normalized to the average grey value for all wildtype mitochondria. A larger value indicates a greater area of the mitochondrion free of dense cristae. Each point represents one mitochondrion.

## References

Abbade, J., Klemetti, M. M., Farrell, A., Ermini, L., Gillmore, T., Sallais, J., Tagliaferro, A., Post, M., & Caniggia, I. (2020). Increased placental mitochondrial fusion in gestational diabetes mellitus: An adaptive mechanism to optimize feto-placental metabolic homeostasis? BMJ Open Diabetes Research and Care, 8(1). 10.1136/bmjdrc-2019-000923

Arnold, F. J., Cui, Y., Michels, S., Colwin, M. R., Stockford, C., Ye, W., Tam, O. H., Menon, S., Situ, W. G., Ehsani, K. C. K., Howard, S., Hammell, M. G., Li, W., & Spada, A. R. La. (2024). TDP-43 dysregulation of polyadenylation site selection is a defining feature of RNA misprocessing in ALS/FTD and related disorders. BioRxiv. 10.1101/2024.01.22.576709

Baughn, M. W., Melamed, Z., López-Erauskin, J., Beccari, M. S., Ling, K., Zuberi, A., Presa, M., Gonzalo-Gil, E., Maimon, R., Vazquez-Sanchez, S., Chaturvedi, S., Bravo-Hernández, M., Taupin, V., Moore, S., Artates, J. W., Acks, E., Ndayambaje, I. S., Agra de Almeida Quadros, A. R., Jafar-Nejad, P., … Cleveland, D. W. (2023). Mechanism of STMN2 cryptic splice-polyadenylation and its correction for TDP-43 proteinopathies. Science (New York, N.Y.), 379(6637), 1140–1149. 10.1126/science.abq5622

Beirowski, B., Gustin, J., Armour, S. M., Yamamoto, H., Viader, A., North, B. J., Michán, S., Baloh, R. H., Golden, J. P., Schmidt, R. E., Sinclair, D. A., Auwerx, J., & Milbrandt, J. (2011). Sir-two-homolog 2 (Sirt2) modulates peripheral myelination through polarity protein Par-3/atypical protein kinase C (aPKC) signaling. Proceedings of the National Academy of Sciences of the United States of America, 108(43), E952–61. 10.1073/pnas.1104969108

Bright, F., Chan, G., van Hummel, A., Ittner, L. M., & Ke, Y. D. (2021). Tdp-43 and inflammation: Implications for amyotrophic lateral sclerosis and frontotemporal dementia. In International Journal of Molecular Sciences (Vol. 22, Issue 15). 10.3390/ijms22157781

Broadhead, M. J., Bonthron, C., Waddington, J., Smith, W. V., Lopez, M. F., Burley, S., Valli, J., Zhu, F., Komiyama, N. H., Smith, C., Grant, S. G. N., & Miles, G. B. (2022). Selective vulnerability of tripartite synapses in amyotrophic lateral sclerosis. Acta Neuropathologica, 143(4). 10.1007/s00401-022-02412-9

Bryce-Smith, S., Brown, A.-L., Mehta, P. R., Mattedi, F., Mikheenko, A., Barattucci, S., Zanovello, M., Dattilo, D., Yome, M., Hill, S. E., Qi, Y. A., Wilkins, O. G., Sun, K., Ryadnov, E., Wan, Y., Consortium, N. A. L. S., Vargas, J. N. S., Birsa, N., Raj, T., … Fratta, P. (2024). TDP-43 loss induces extensive cryptic polyadenylation in ALS/FTD. BioRxiv. 10.1101/2024.01.22.576625

Chaplan, S. R., Bach, F. W., Pogrel, J. W., Chung, J. M., & Yaksh, T. L. (1994). Quantitative assessment of tactile allodynia in the rat paw. Journal of Neuroscience Methods, 53(1), 55–63. 10.1016/0165-0270(94)90144-9

Dixon, W. J. (1980). Efficient analysis of experimental observations. Annual Review of Pharmacology and Toxicology, 20. 10.1146/annurev.pa.20.040180.002301

Geisler, S., Doan, R. A., Strickland, A., Huang, X., Milbrandt, J., & DiAntonio, A. (2016). Prevention of vincristine-induced peripheral neuropathy by genetic deletion of SARM1 in mice. Brain, 139(Pt 12), 3092–3108. 10.1093/brain/aww251

Graf, E. R., Heerssen, H. M., Wright, C. M., Davis, G. W., & di Antonio, A. (2011). Stathmin is required for stability of the drosophila neuromuscular junction. Journal of Neuroscience, 31(42). 10.1523/JNEUROSCI.2024-11.2011

Guerra San Juan, I., Nash, L. A., Smith, K. S., Leyton-Jaimes, M. F., Qian, M., Klim, J. R., Limone, F., Dorr, A. B., Couto, A., Pintacuda, G., Joseph, B. J., Whisenant, D. E., Noble, C., Melnik, V., Potter, D., Holmes, A., Burberry, A., Verhage, M., & Eggan, K. (2022). Loss of mouse Stmn2 function causes motor neuropathy. Neuron. 10.1016/j.neuron.2022.02.011

Guerrero, E. N., Mitra, J., Wang, H., Rangaswamy, S., Hegde, P. M., Basu, P., Rao, K. S., & Hegde, M. L. (2019). Amyotrophic lateral sclerosis-associated TDP-43 mutation Q331K prevents nuclear translocation of XRCC4-DNA ligase 4 complex and is linked to genome damage-mediated neuronal apoptosis. Human Molecular Genetics, 28(18), 3161–3162. 10.1093/hmg/ddz141

Jadiya, P., Garbincius, J. F., & Elrod, J. W. (2021). Reappraisal of metabolic dysfunction in neurodegeneration: Focus on mitochondrial function and calcium signaling. In Acta Neuropathologica Communications (Vol. 9, Issue 1). 10.1186/s40478-021-01224-4

Jeong, Y. H., Ling, J. P., Lin, S. Z., Donde, A. N., Braunstein, K. E., Majounie, E., Traynor, B. J., LaClair, K. D., Lloyd, T. E., & Wong, P. C. (2017). Tdp-43 cryptic exons are highly variable between cell types. Molecular Neurodegeneration. 10.1186/s13024-016-0144-x

Klim, J. R., Williams, L. A., Limone, F., Guerra San Juan, I., Davis-Dusenbery, B. N., Mordes, D. A., Burberry, A., Steinbaugh, M. J., Gamage, K. K., Kirchner, R., Moccia, R., Cassel, S. H., Chen, K., Wainger, B. J., Woolf, C. J., & Eggan, K. (2019). ALS-implicated protein TDP-43 sustains levels of STMN2, a mediator of motor neuron growth and repair. Nature Neuroscience. 10.1038/s41593-018-0300-4

Krus, K. L., Strickland, A., Yamada, Y., Devault, L., Schmidt, R. E., Bloom, A. J., Milbrandt, J., & DiAntonio, A. (2022). Loss of Stathmin-2, a hallmark of TDP-43-associated ALS, causes motor neuropathy. BioRxiv. 10.1101/2022.03.13.484188

Li, Y., Tian, Y., Pei, X., Zheng, P., Miao, L., Li, L., Luo, C., Zhang, P., Jiang, B., Teng, J., Huang, N., & Chen, J. (2023). SCG10 is required for peripheral axon maintenance and regeneration in mice. Journal of Cell Science, 136(12). 10.1242/jcs.260490

Ling, J. P., Pletnikova, O., Troncoso, J. C., & Wong, P. C. (2015). TDP-43 repression of nonconserved cryptic exons is compromised in ALS-FTD. Science. 10.1126/science.aab0983

Ling, S. C., Polymenidou, M., & Cleveland, D. W. (2013). Converging mechanisms in als and FTD: Disrupted RNA and protein homeostasis. In Neuron. 10.1016/j.neuron.2013.07.033

López-Erauskin, J., Bravo-Hernandez, M., Presa, M., Baughn, M. W., Melamed, Z., Beccari, M. S., Agra de Almeida Quadros, A. R., Arnold-Garcia, O., Zuberi, A., Ling, K., Platoshyn, O., Niño-Jara, E., Ndayambaje, I. S., McAlonis-Downes, M., Cabrera, L., Artates, J. W., Ryan, J., Hermann, A., Ravits, J., … Lagier-Tourenne, C. (2024). Stathmin-2 loss leads to neurofilament-dependent axonal collapse driving motor and sensory denervation. Nature Neuroscience, 27(1). 10.1038/s41593-023-01496-0

Ma, X. R., Prudencio, M., Koike, Y., Vatsavayai, S. C., Kim, G., Harbinski, F., Briner, A., Rodriguez, C. M., Guo, C., Akiyama, T., Schmidt, H. B., Cummings, B. B., Wyatt, D. W., Kurylo, K., Miller, G., Mekhoubad, S., Sallee, N., Mekonnen, G., Ganser, L., … Gitler, A. D. (2022). TDP-43 represses cryptic exon inclusion in the FTD-ALS gene UNC13A. Nature, 603(7899), 124–130. 10.1038/s41586-022-04424-7

Melamed, Z., López-Erauskin, J., Baughn, M. W., Zhang, O., Drenner, K., Sun, Y., Freyermuth, F., McMahon, M. A., Beccari, M. S., Artates, J. W., Ohkubo, T., Rodriguez, M., Lin, N., Wu, D., Bennett, C. F., Rigo, F., Da Cruz, S., Ravits, J., Lagier-Tourenne, C., & Cleveland, D. W. (2019). Premature polyadenylation-mediated loss of stathmin-2 is a hallmark of TDP-43-dependent neurodegeneration. Nature Neuroscience. 10.1038/s41593-018-0293-z

Morii, H., Shiraishi-Yamaguchi, Y., & Mori, N. (2006). SCG10, a microtubule destabilizing factor, stimulates the neurite outgrowth by modulating microtubule dynamics in rat hippocampal primary cultured neurons. Journal of Neurobiology. 10.1002/neu.20295

National Institute of Health. (2022). A Study Evaluating the Safety and Tolerability of QRL-201 in ALS (https://Clinicaltrials.gov Identifier NCT05633459). Retrieved from https://www.clinicaltrials.gov/study/NCT05633459.

Neumann, M., Sampathu, D. M., Kwong, L. K., Truax, A. C., Micsenyi, M. C., Chou, T. T., Bruce, J., Schuck, T., Grossman, M., Clark, C. M., McCluskey, L. F., Miller, B. L., Masliah, E., Mackenzie, I. R., Feldman, H., Feiden, W., Kretzschmar, H. A., Trojanowski, J. Q., & Lee, V. M. Y. (2006). Ubiquitinated TDP-43 in frontotemporal lobar degeneration and amyotrophic lateral sclerosis. Science. 10.1126/science.1134108

Sasaki, Y., Hackett, A. R., Kim, S., Strickland, A., & Milbrandt, J. (2018). Dysregulation of NAD+ metabolism induces a Schwann cell dedifferentiation program. Journal of Neuroscience. 10.1523/JNEUROSCI.3304-17.2018

Shin, J. E., Geisler, S., & DiAntonio, A. (2014). Dynamic regulation of SCG10 in regenerating axons after injury. Experimental Neurology, 252. 10.1016/j.expneurol.2013.11.007

Shin, J. E., Miller, B. R., Babetto, E., Cho, Y., Sasaki, Y., Qayum, S., Russler, E. V., Cavalli, V., Milbrandt, J., & DiAntonio, A. (2012). SCG10 is a JNK target in the axonal degeneration pathway. Proceedings of the National Academy of Sciences of the United States of America. 10.1073/pnas.1216204109

Sreedharan, J., Blair, I. P., Tripathi, V. B., Hu, X., Vance, C., Rogelj, B., Ackerley, S., Durnall, J. C., Williams, K. L., Buratti, E., Baralle, F., De Belleroche, J., Mitchell, J. D., Leigh, P. N., Al-Chalabi, A., Miller, C. C., Nicholson, G., & Shaw, C. E. (2008). TDP-43 mutations in familial and sporadic amyotrophic lateral sclerosis. Science, 319(5870). 10.1126/science.1154584

van Es, M. A., Hardiman, O., Chio, A., Al-Chalabi, A., Pasterkamp, R. J., Veldink, J. H., & van den Berg, L. H. (2017). Amyotrophic lateral sclerosis. The Lancet, 390(10107), 2084–2098. 10.1016/S0140-6736(17)31287-4

Wang, W., Arakawa, H., Wang, L., Okolo, O., Siedlak, S. L., Jiang, Y., Gao, J., Xie, F., Petersen, R. B., & Wang, X. (2017). Motor-Coordinative and Cognitive Dysfunction Caused by Mutant TDP-43 Could Be Reversed by Inhibiting Its Mitochondrial Localization. Molecular Therapy,_j: The Journal of the American Society of Gene Therapy, 25(1), 127– 139. 10.1016/j.ymthe.2016.10.013

Wang, W., Li, L., Lin, W. L., Dickson, D. W., Petrucelli, L., Zhang, T., & Wang, X. (2013). The ALS disease-associated mutant TDP-43 impairs mitochondrial dynamics and function in motor neurons. Human Molecular Genetics, 22(23). 10.1093/hmg/ddt319

Wang, W., Wang, L., Lu, J., Siedlak, S. L., Fujioka, H., Liang, J., Jiang, S., Ma, X., Jiang, Z., da Rocha, E. L., Sheng, M., Choi, H., Lerou, P. H., Li, H., & Wang, X. (2016). The inhibition of TDP-43 mitochondrial localization blocks its neuronal toxicity. Nature Medicine, 22(8), 869– 878. 10.1038/nm.4130

White, M. A., Kim, E., Duffy, A., Adalbert, R., Phillips, B. U., Peters, O. M., Stephenson, J., Yang, S., Massenzio, F., Lin, Z., Andrews, S., Segonds-Pichon, A., Metterville, J., Saksida, L. M., Mead, R., Ribchester, R. R., Barhomi, Y., Serre, T., Coleman, M. P., … Sreedharan, J. (2018). TDP-43 gains function due to perturbed autoregulation in a Tardbp knock-in mouse model of ALS-FTD. Nature Neuroscience. 10.1038/s41593-018-0113-5

Yu, C. H., Davidson, S., Harapas, C. R., Hilton, J. B., Mlodzianoski, M. J., Laohamonthonkul, P., Louis, C., Low, R. R. J., Moecking, J., De Nardo, D., Balka, K. R., Calleja, D. J., Moghaddas, F., Ni, E., McLean, C. A., Samson, A. L., Tyebji, S., Tonkin, C. J., Bye, C. R., … Masters, S. L. (2020). TDP-43 Triggers Mitochondrial DNA Release via mPTP to Activate cGAS/STING in ALS. Cell, 183(3). 10.1016/j.cell.2020.09.020

Zeng, Y., Lovchykova, A., Akiyama, T., Liu, C., Guo, C., Jawahar, V. M., Sianto, O., Calliari, A., Prudencio, M., Dickson, D. W., Petrucelli, L., & Gitler, A. D. (2024). TDP-43 nuclear loss in FTD/ALS causes widespread alternative polyadenylation changes. BioRxiv. 10.1101/2024.01.22.575730

