## Supplementary figures and images for "Reduced STMN2 and pathogenic TDP-43, two hallmarks of ALS, synergize to accelerate motor decline in mice"

### Supplemental Figure 1

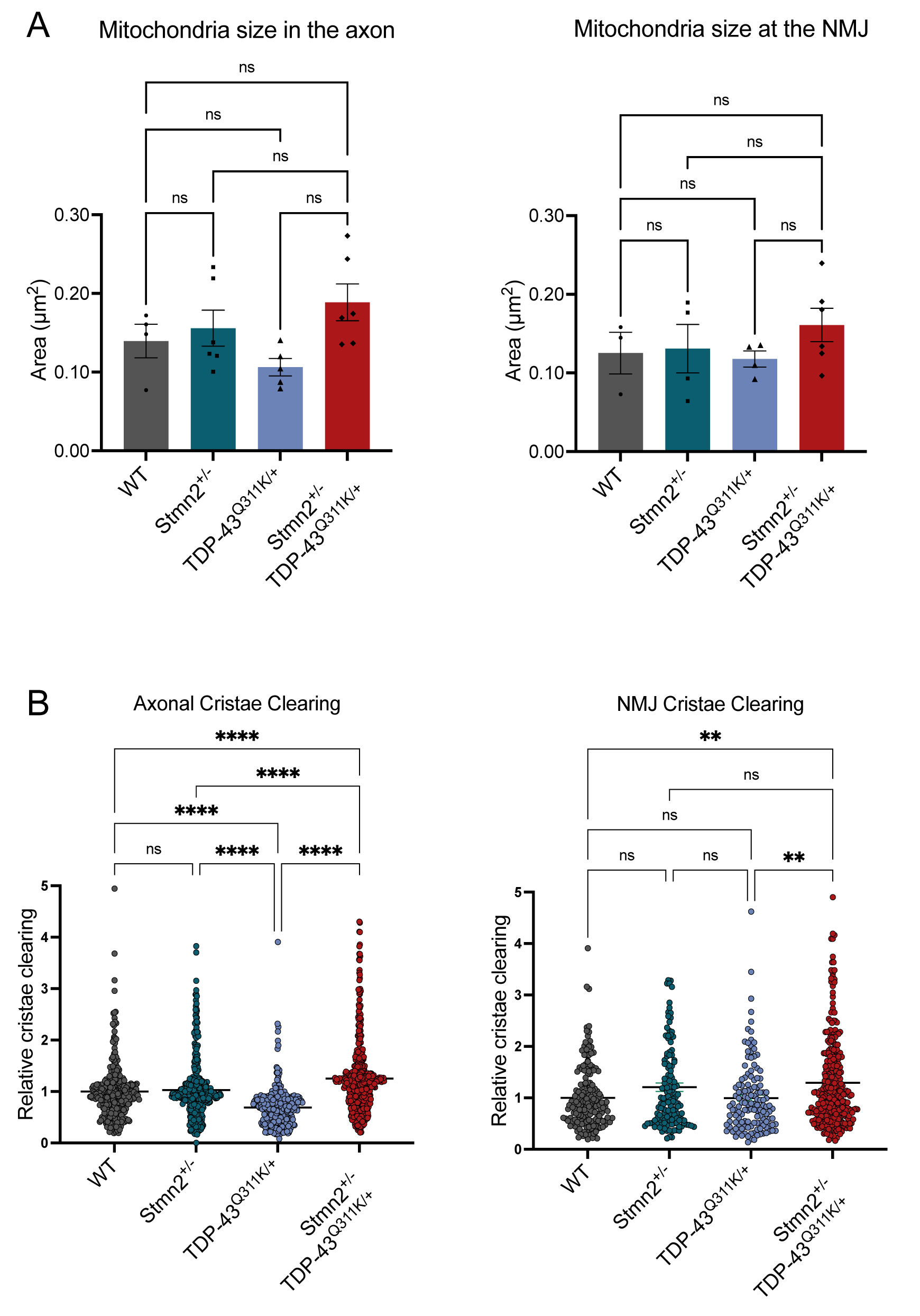
